# Variation in the geometry of concept manifolds across human visual cortex

**DOI:** 10.1101/2024.11.26.625280

**Authors:** Ghislain St-Yves, Kendrick Kay, Thomas Naselaris

## Abstract

Brain activity patterns in high-level visual cortex support accurate linear classification of visual concepts (e.g., objects or scenes). It has long been appreciated that the accuracy of linear classification in any brain area depends on the geometry of its concept manifolds—sets of brain activity patterns that encode images of a concept. However, it is unclear how the geometry of concept manifolds differs between regions of visual cortex that support accurate classification and those that don’t, or how it differs between visual cortex and deep neural networks (DNNs). We estimated geometric properties of concept manifolds that, per a recent theory, directly determine the accuracy of simple “few-shot” linear classifiers. Using a large fMRI dataset, we show that variation in classification accuracy across human visual cortex is driven by a variation in a single geometric property: the distance between manifold centers (“geometric Signal”). In contrast, variation in classification accuracy across most DNN layers is driven by an increase in the effective number of manifold dimensions (“Dimensionality”). Despite this difference in the geometric properties that affect few-shot classification performance in the brain and DNNs, we find that Signal and Dimensionality are strongly, negatively correlated: when Signal increases across brain regions or DNN layers, Dimensionality decreases, and vice versa. We conclude that visual cortex and DNNs deploy different geometric strategies for accurate linear classification of concepts, even though both are subject to the same constraint.

## Introduction

Visual representations in the human brain must support the accurate classification of visual concepts that pertain to objects or scenes. It is currently unclear what distinguishes image representations in cortical regions where linear read-outs of brain activity patterns accurately classify visual concepts from representations in cortical regions where linear read-out fails. Although Deep Neural Networks (DNNs) trained to classify objects provide some compelling clues Yamins et al. [2014], Khaligh-Razavi and Kriegeskorte [2014], Richards et al. [2019], solutions to the problem of concept classification encoded in the activity pattern of DNNs and the human brain can differ in important ways Geirhos et al. [2018], Jacob et al. [2021].

We define a visual concept as any set of images that share some attribute (e.g., the set of all images that depict a dog). We refer to images in the set as “concept exemplars”. If we take the neural representation of an image to be a point in the space of brain activity (or digital activity, in the case of DNNs), then the neural representation of a concept is the set of points that encode its exemplars. Such sets of points in brain activity space have been referred to as “concept manifolds” Sorscher et al. [2022]. It has long been appreciated that accurate classification of visual concepts via linear read-out is possible only when concept manifolds are “disentangled” DiCarlo and Cox [2007]. Intuitively, disentangled manifolds occupy different, non-overlapping regions of brain space. According to this intuition, we can understand how image representations in some parts of the human brain support concept classification by identifying the geometric properties of their concept manifolds that make them more or less disentangled. The geometry of concept manifolds is currently a topic of intense investigation Jazayeri and Ostojic [2021], Chung and Abbott [2021], Li et al. [2024].

Recently, Sorscher et al. Sorscher et al. [2022] developed a theory that relates a set of intuitive and relatively easy-to-measure geometric properties to the error of so-called “few-shot” linear concept classifiers, i.e., linear classifiers that are learned using a small number of training examples. The theory thus provides a way to identify and estimate from brain activity the specific geometric properties that make concept manifolds more disentangled in some cortical regions than in others.

Importantly, the theory reveals that a change in classification accuracy between cortical regions can be effected by many different kinds of changes to manifold geometry. For example, according to the theory a higher-level cortical region could increase classification accuracy relative to V1 by increasing the distance between manifold centers, by increasing the dimensionality of manifolds, by rotating manifolds such that their primary axes of variation are less aligned, or by effecting some combination of these changes. It is currently unknown which of these possible “geometric strategies” is most evident in human brain activity patterns measured with fMRI across the entire visual cortex, nor how strategies inferred from analyzing human fMRI data might differ from those of DNNs.

In this work, we utilize the theory of Sorscher et al. and brain activity patterns in the Natural Scenes Dataset Allen et al. [2022] to estimate the variation in geometric properties of concept manifolds across sequences of human visual cortical areas that are ordered by decreasing classification error. We identify specific geometric properties that are primarily responsible for the decrease in few-shot classification error across visual cortical areas, and reveal how much of the overall variation in geometry increases, decreases, or has no impact on variation in classification accuracy. We also contrast the variation in the geometry of concept manifolds across human visual cortex to variation in the geometry of concept manifolds across the layers of deep neural networks optimized to perform many-shot classification over thousands of object categories.

## Results

### Concept manifold geometry and few-shot classification error

Consider two concepts, *a* and *b*. A key insight of the theory of Sorscher et al. is that even though the manifolds for concepts *a* and *b* may have quite complex structure, the error of a linear few-shot classifier that is trained to discriminate between the concepts depends upon a small number of intuitive geometric properties: the distance between the centers of the manifolds (“geometric Signal”); the radius of manifold *a* relative to the radius of manifold *b* (“Bias”); the number of independent directions of variation that are utilized by a manifold (“Dimensionality”); the alignment of principal manifold directions with each other (Noise-Noise Overlap); and the alignment of the principal manifold directions with the Signal vector that connects the centers of the manifolds (Noise-Signal Overlap).

Sorscher et al. introduced a critical quantity, the *geometric Signal-to-Noise ratio* (SNR(*m*)), that relates all of these geometric properties to *m* -shot classification error:

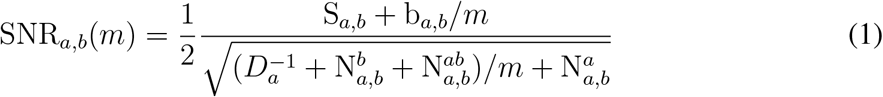

where S_*a,b*_ and b_*a,b*_ are the geometric Signal and Bias terms respectively. The denominator consists of all the nuisance factors: the Noise-Signal Overlap 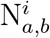 for manifold *i* and the NoiseNoise Overlap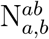 that capture the overlap between all of the manifolds’ directions (Fig. 1A).

**Figure 1:**
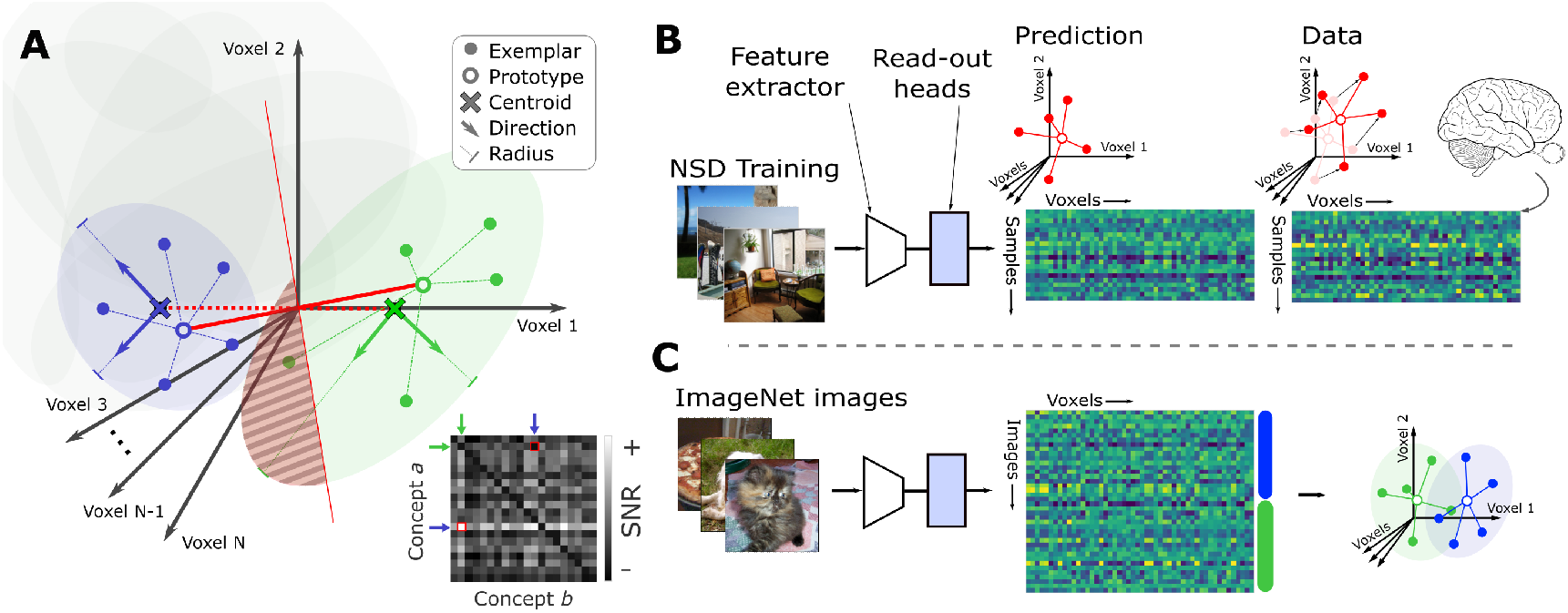
Measuring geometric properties of concept manifolds in the space of human brain activity. **(A)** Concept manifolds (shown here as ellipses) for two concepts (blue and green, respectively) in the space of brain activity (each axis of the space corresponds to the activity in a single voxel). To construct an *m*-shot (*m* = 5 in this illustration) classifier, prototypes (open circles) are estimated by computing the mean of *m* exemplar activity patterns (filled circles). At test, activity patterns are projected onto the vector (thick red line) connecting the concept prototypes, and thresholded via distance to the mid-point (thin red line) between prototypes. According to the theory of Sorcher et al., the error of such an *m*-shot classifier is determined by several geometric properties of the concept manifolds: geometric Signal is distance (dashed red line) between the true manifold centroids (dark x-marks) relative to the radii (lines with feet) of the principal directions (thick green and blue arrows); Bias is the relative radius of each manifold; Noise-Noise Overlap measures the alignment of the two manifolds’ principal directions; Noise-Signal Overlap is the alignment of the manifolds’ principal directions and the line (dashed red) connecting the centroids; Dimensionality is a measure of how uniformly the variance in concept exemplars is distributed across the axes of the brain space. Collectively, these properties determine a quantity called the “geometric SNR” (inset grayscale matrix) that, per the theory, predicts *m*-shot error for any pair of concepts. In this illustration the *m*-shot error will be non-zero, since at least some of the exemplars (in the striped area) will be mis-classified. **(B)** Inferring geometric properties from fMRI data. Geometric properties and few-shot error are inferred from activity patterns (colored matrices), shown as exemplars (filled circles) and prototypes (open circles) in the space of brain activity measured using fMRI (right; pink) or output from an encoding model (left; red). In our case the encoding model is a deep neural network consisting of a feature extractor (trapezoid) and voxel-wise read-out heads (rectangle) and trained end-to-end to predict brain activity. **(C)** Once trained, an encoding model can be used to generate estimates of activity patterns (matrix) for exemplars (rows) of any pair of visual object categories (blue / green bars) in the ImageNet database. From these, geometric properties of pairs of concept manifolds (green/blue ellipses) are computed.

For all elements, the subscript indices indicate that the element describes the property of concept *a* relative to concept *b*. These quantities are generally asymmetrical with respect to the concept manifolds, meaning it is possible that SNR_*a,b*_(*m*) ≠ SNR_*b,a*_(*m*). The *m*-shot error rate is related to the SNR(*m*) by a Gaussian tail function (dashed black curves in Fig. 4). Finally, *D*_*a*_ is the participation ratio dimension (referred to here as “Dimensionality”) of concept *a*.

As the equation shows, classification error will be small when geometric Signal and Dimensionality are large and Noise-Noise and Noise-Signal Overlap are small. In other words, visual representations support accurate few-shot classification of concepts *a* and *b* when their manifolds are far apart, when the concepts utilize multiple directions of variation, and when the principal directions of the manifolds do not align with each other or with the Signal vector.

It is important to note that the expression for geometric SNR does not reveal any explicit dependencies between the geometric properties. Thus, as far as the expression for geometric SNR is concerned, there are many different patterns of co-variation in geometric properties across visual cortex that can induce an increase or decrease in SNR. Variation in geometric SNR could emerge as a result of “coordinated” co-variation, in which variation in all of the constituent geometric properties affect geometric SNR in the same way, or as a result of “antagonistic” covariation, in which variation in the constituent properties have opposing effects on geometric SNR (as seen in the example below; Fig. 2B). Also consistent with the theory is an interesting special case of antagonistic co-variation, in which variation in all geometric properties cancel each others’ effects, resulting in potentially profound variation in manifold geometry across visual cortex, but no variation in geometric SNR. From the limited perspective of understanding concept classification, such a profile of co-variation in geometric properties could be considered “non-functional”.

**Figure 2:**
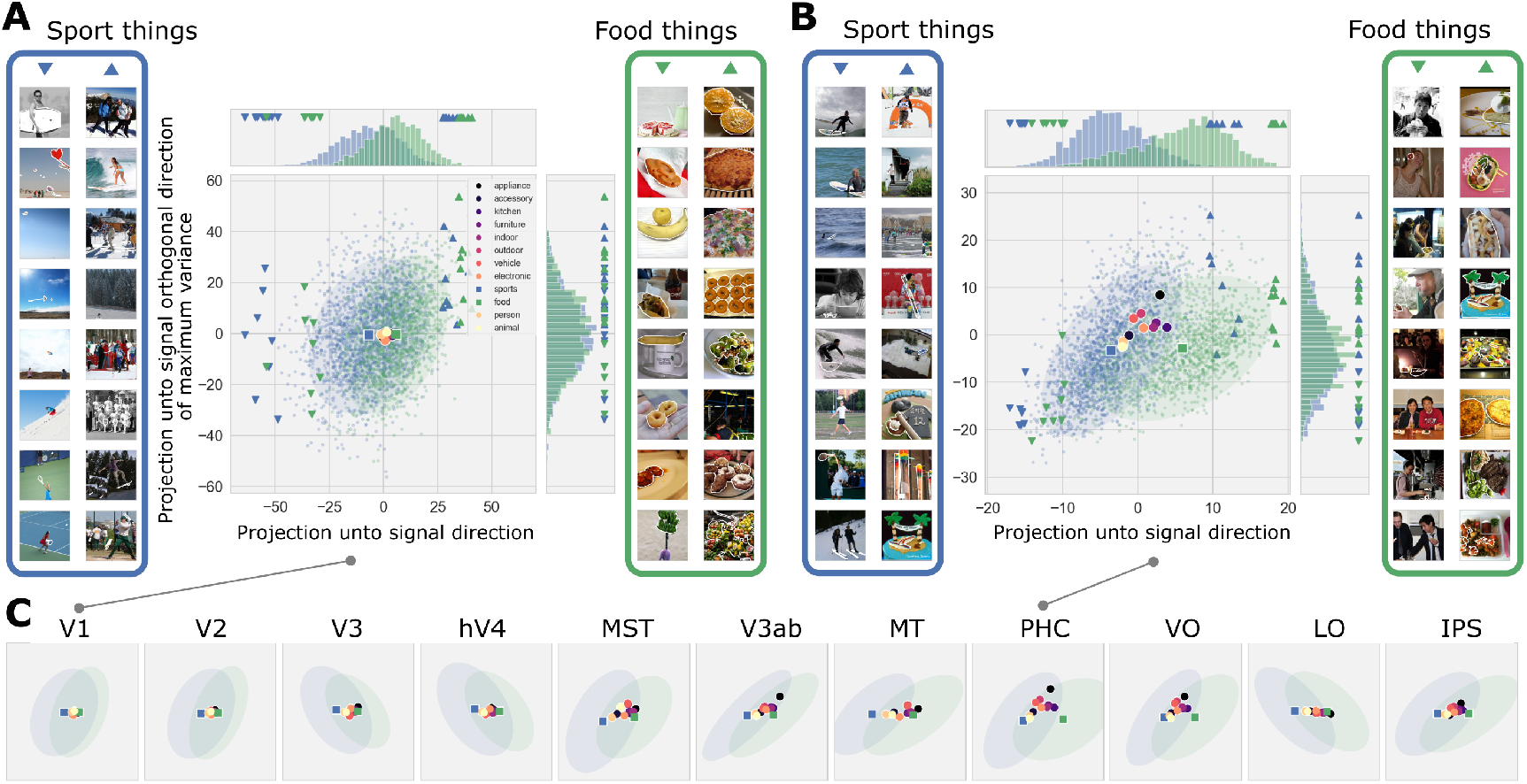
Illustration of two concept manifolds for brain regions of interest across human visual cortex. Projections of voxel activity patterns in **(A)** V1 and **(B)** PHC corresponding to two concepts: “sport”-related things (each blue dot is one activity pattern) and “food”-related things (green dots). Activity patterns are projected onto the Signal direction (x-axis), i.e., the vector connecting the centroids (squares) of each concept manifold, and the direction orthogonal to the signal direction (y-axis) with maximum variance for the two concepts. The centroids for other concepts (large circles) are displayed for reference. Images (panels to left and right) associated with extremal activity patterns (triangles) indicate some of the image features that vary along the signal direction. **(C)** Summaries of sport-things and food-things manifolds (blue and green ellipses, resp.; axes same as in A,B) for all brain ROIs considered here.

To gain an intuition for how these various geometric quantities might vary across human visual cortex, we visualized 2D projections of concept manifolds using brain activity patterns in the Natural Scenes Dataset (NSD), a massive sampling of blood-oxygenation-level-dependent (BOLD) activity in eight subjects using ultra-high field fMRI (7T, 1.8-mm resolution). Subjects each viewed 9000–10,000 natural scenes (sampled from the Microsoft Common Objects in Context database Lin et al. [2015]) presented (3-s exposure) repeatedly (three times typically), yielding 22–30K trials for individual subjects and a total of 213K trials across subjects.

We visualized manifolds for the concepts “has something to do with sports” and “has some-thing to do with food” in visual regions of interest (ROIs; V1, V2, V3, V4, V3a, V3b, lateral occipital complex (LOC), ventral occipital cortex (VO), intraparietal sulcus (IPS), middle temporal area (MT), medial superior temporal area (MST) and peri-hippocampal gyrus (PHC)) that collectively span most of visually-responsive cortex (Fig. 2). As expected, the concept mani-folds in higher areas appeared to be more disentangled than concept manifolds in lower visual areas, in the sense that they overlap less in the 2D plane used here for visualization (Fig. 2). The visualizations offered some clues about what particular geometric properties make the manifolds for these specific concepts more disentangled in higher than in lower visual areas. For example, in PHC the manifold centers are farther apart (relative to their respective radii) than in V1, suggesting that manifolds in PHC are more disentangled than in V1 due to an increase in geometric Signal.

Interestingly, in PHC the “sport things” manifold was compressed towards a principal direction that was clearly rotated toward the Signal vector, indicating a decrease in Dimensionality and an increase in noise-signal Overlap in PHC relative to V1. Per the theory of Sorscher et al., linear classification error increases as Dimensionality decreases, and as Noise-Signal Overlap increases; thus, manifolds in PHC were more disentangled than in V1 in spite of a relative decrease in Dimensionality and increase in Noise-Signal Overlap. This is an illustration of “antagonistic” co-variation because a change in one geometric property (Signal) effected an increase in SNR, while a change other geometric properties (Dimensionality and Overlap) mitigated the effect.

Having established these intuitions, we used the NSD to conduct a systematic exploration of few-shot classification error and manifold geometry across visual cortex.

### Geometric properties of concept manifolds precisely determine the error of few-shot classifiers estimated from both human brain activity patterns and the outputs of an encoding model

Before estimating concept manifold geometry, we first empirically measured how the error of few-shot classifiers for thousands of visual concepts varied across the visual cortex. To control for the effects of inter-trial variability (i.e., the variability of activity in response to multiple presentations of the same stimulus), and to expand the concept repertoire, we first investigated synthetic brain activity patterns output by a brain-optimized encoding model. The encoding model, known as GNet St-Yves et al. [2023], is a deep neural network that was trained end-to-end using measured fMRI brain activity patterns in the NSD. GNet outputs a predicted brain activity pattern in response to each input image. Using GNet, we were able to construct fewshot classifiers from noiseless predicted brain activity patterns for roughly one million concept pairs from the ImageNet database. For each region of interest (ROI) and pair of concepts, we resampled exemplars to construct multiple 5-shot linear classifiers, computed their errors (the fraction of incorrect classifications on a held-out test sample), and then averaged errors across the classifiers.

In each ROI, error decreased with the semantic distance (number of “hops” between concepts on the WordNet graph Fellbaum [2010]) between the concept pairs (Fig. 3A). Across ROIs, error induced an unsurprising posterior to anterior ordering that was consistent across most levels of semantic distance (Fig. 3B). Under the ordering induced by the largest semantic distance (darkest curve in Fig. 3B) classification error decreases rapidly across early and intermediate visual areas V1–V4, then more subtly across a collection of more anterior lateral (MST, MT, LO), dorsal (V3AB) and ventral (PHC, VO) ROIs, and terminates with an ROI spanning much of the intraparietal sulcus (IPS), suggesting that much of the geometric variation that supports accurate linear classification in high-level areas occurs across early and intermediate visual areas.

**Figure 3:**
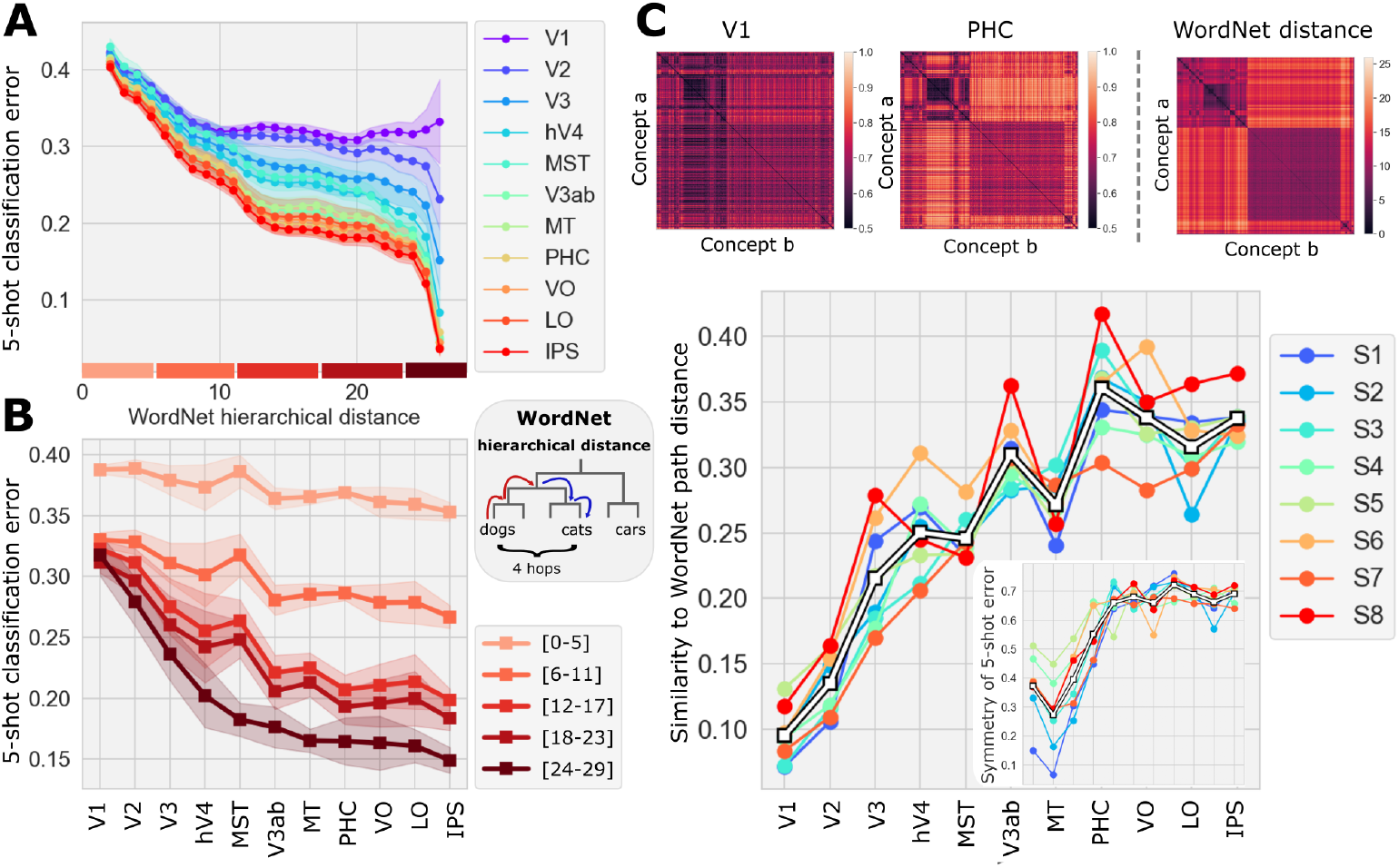
Variation in few-shot classification error across human visual cortex. **(A)** Average 5-shot error (1 - fraction of correctly classified exemplars; y-axis) as a function of WordNet path distance (x-axis; see inset) between the concept pairs in all ROIs (colored curves) considered here. **(B)** 5-shot error across ROIs (x-axis) for pairs of concepts across a range of WordNet distances (colors). **(C)** Top left: 5-shot success (1 - error; color bar) matrices for V1 and PHC. Top right: WordNet distance matrix for all concept pairs. Bottom: Correlation (y-axis) between 5-shot success and WordNet path distance for all pairs of concepts in ImageNet across ROIs (x-axis). The inset shows a measure of the symmetry of the 5-shot error matrices.

As error decreased across this sequence, the concept-by-concept matrix of classification accuracy (1 - error) increasingly resembled concept-by-concept semantic distances in WordNet (Fig. 3C). Note that although semantic distances are symmetric on the WordNet graph, classification errors need not be. For example, a classifier that discriminates between “dog” and “cat” may correctly identify “dog” exemplars more often than it correctly identifies “cat” exemplars. Interestingly, classification error matrices became more symmetric with progression along the sequence of ROIs (Fig. 3C, inset), and the similarity between few-shot accuracy and hierarchical distance continues to increase even after symmetry plateaus. This suggests that the increase in alignment between 5-shot error and WordNet distance is not fully explained by an increase in the symmetry of 5-shot errors alone.

Having calculated the average few-shot classification error for each ROI, we then confirmed that the expression for geometric SNR (Equation 1) correctly predicted classification error in our particular setting—i.e., concept manifolds that are sets of brain activity patterns measured in the human visual cortex using fMRI. We first confirmed the theory for measured brain activity patterns. To do this, we divided images in the NSD into 12 super-category concepts (see legend in 2A). For each concept, we selected 580 activity patterns and used these to calculate, for each pair of concepts, the geometric SNR as specified by Equation 1. We found that geometric SNR was an extremely accurate predictor of 5-shot error in our setting (Fig. 4A, left and right).

**Figure 4:**
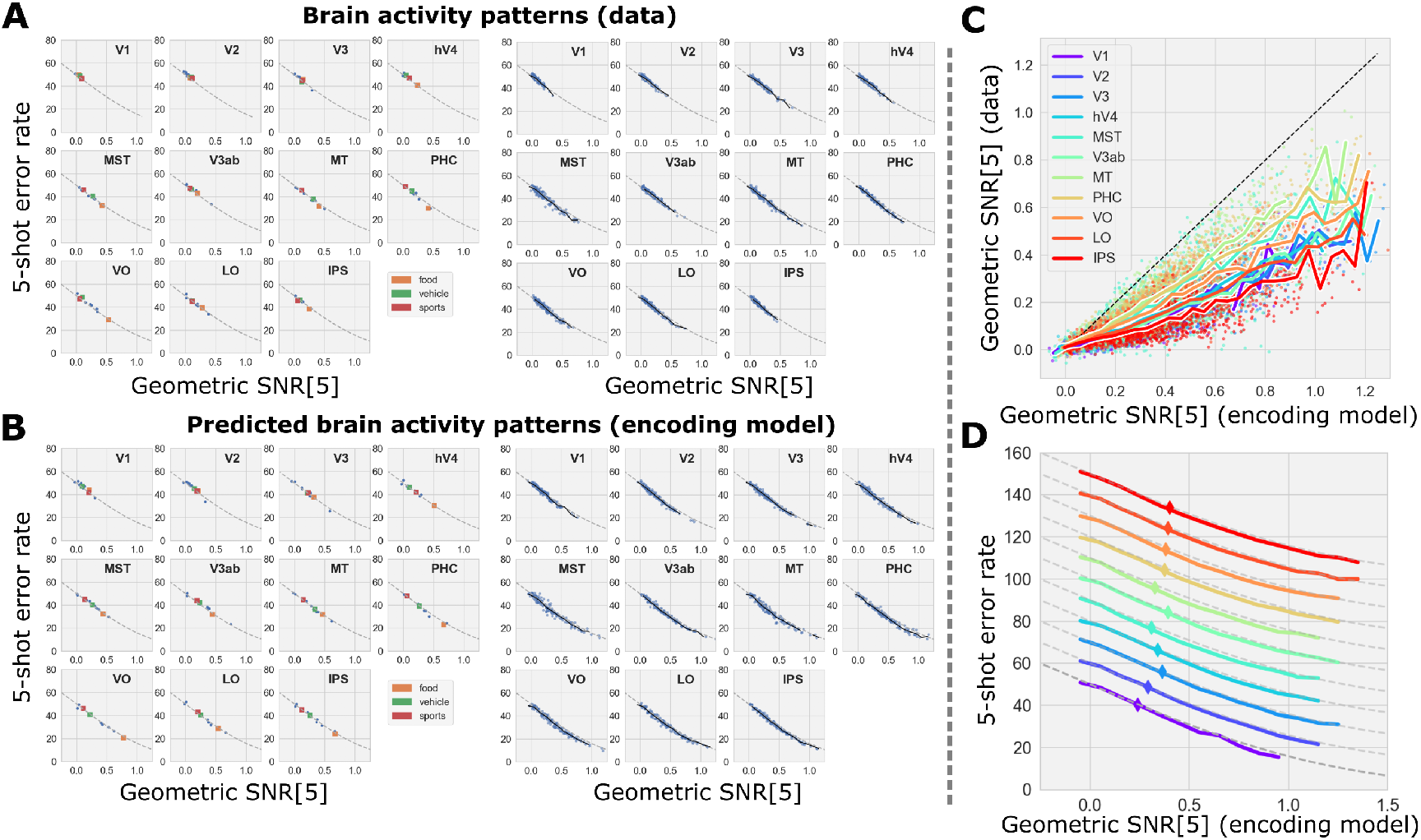
Predicting few-shot classification error from geometric properties of concept manifolds. **(A)** (left) Theoretical relationship (dashed curve) between 5-shot error (y-axis) and geometric SNR (x-axis) compared to empirical measurements of each for pairs of concepts that include “person” and “food” (orange), “vehicle” (green), “sport” (red), and all other categories (blue) calculated directly for brain activity patterns (dots) in response to presentation of images containing instances of the various concepts. (right) All pairs of concepts listed in Fig. 2A. **(B)** Same format as A where the geometric SNR and 5-shot error has been calculated using synthetic brain activity patterns output by an encoding model trained to predict brain activity in visual cortex. **(C)** SNR for pairs of concepts using activity predicted by the encoding model (x-axis) and measured by fMRI (y-axis) across all ROIs (colored curves). **(D)** Binned average of empirical 5-shot error (x-axis) as a function of average geometric SNR per the theory of Sorcher et al. (dashed lines) and for each ROI (colors). Curves have been shifted upward by 10% in order to disambiguate them. The diamond symbol shows the mean 5-shot error and mean SNR[5] over all concept pairs.

We then confirmed the theory for synthetic brain activity patterns output by the GNet encoding model and found that, again, geometric SNR predicted the variation in 5-shot error across many pairs of concepts. In all areas, geometric SNR derived from activity patterns output by our encoding model was strongly correlated with–but much higher than–geometric SNR derived from measured activity (Fig. 4C). The increase in geometric SNR for the encoding model outputs is due to the fact that the encoding model eliminates inter-trial variability in the brain activity patterns, and thus isolates the semantic information they encode. For this reason, we based subsequent analyses on the outputs of our encoding model. Finally, in what follows we will often consider the average of the classification error and other geometric properties across many pairs of concepts within a single ROI; thus, we confirmed that averaged geometric SNR predicted the average classification error (Fig. 4D).

### Variation in few-shot classification accuracy across visual cortex depends primarily upon variation in geometric Signal

Having confirmed the predictive validity of the expression for geometric SNR in our particular setting, we next studied the profile of variation in geometric SNR and all other geometric properties identified by the theory of Sorscher et al. across our brain ROIs (Fig. 5). For each ROI, we averaged geometric SNR across pairs of ImageNet concepts. The ordering induced by average geometric SNR was identical to the ordering induced by 5-shot classification error from concepts separated by more than 23 “hops” on the WordNet graph. We also averaged geometric SNR across pairs of COCO super-categories, inducing a largely consistent (although not identical) ordering of ROIs.

**Figure 5:**
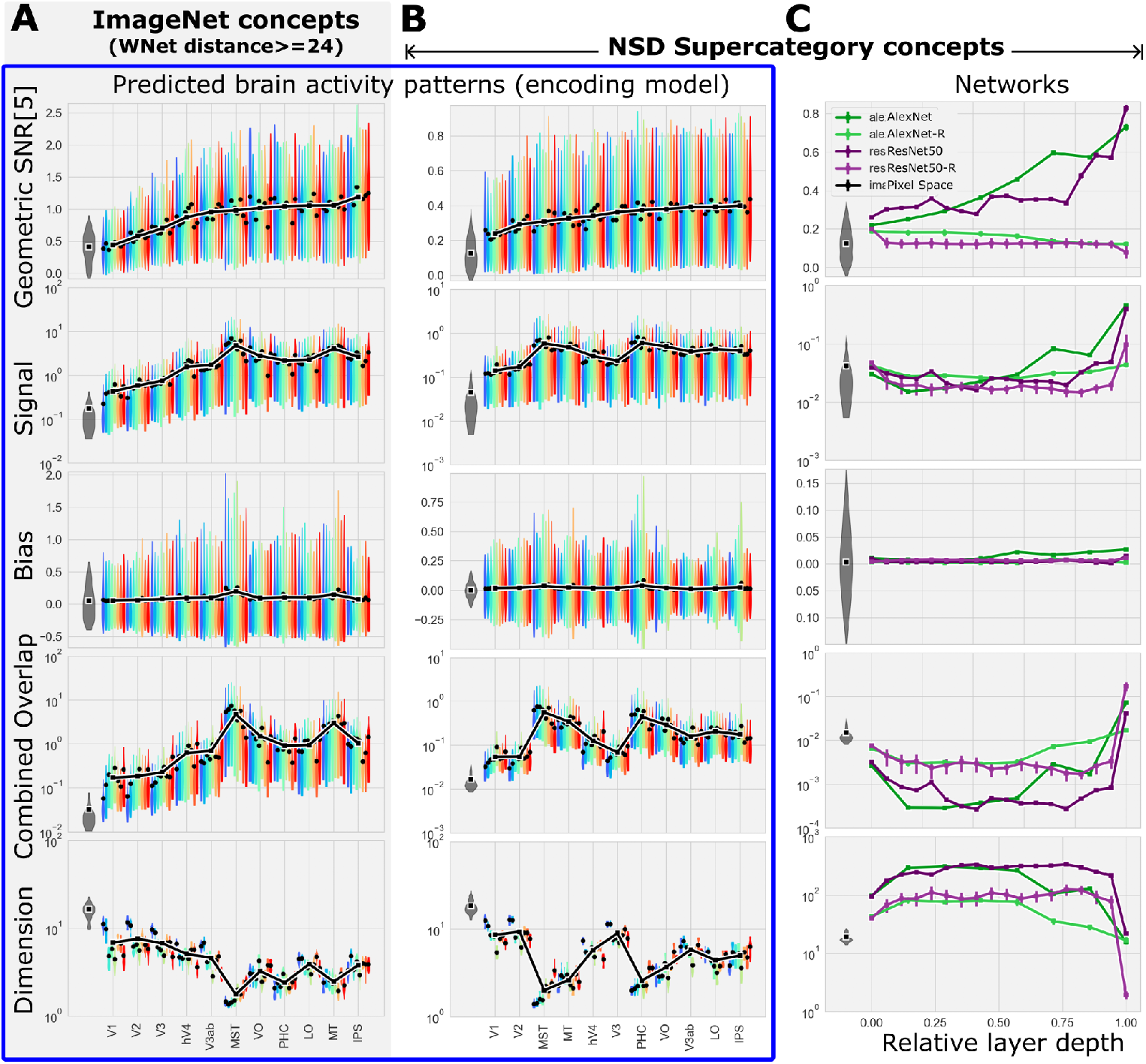
Variation in the geometry of concept manifolds across human visual cortex and layers of deep neural networks. **(A)** The average (black dots) geometric SNR, Signal, Bias, combined Overlap and Dimensionality for each subject (colors; thick black curve is cross-subject average; maximum width of each violin shape indicates modal value, extrema indicate 5th and 95th percentiles) estimated from predicted brain activity patterns and 1000 ImageNet categories for all ROIs ordered by ascending SNR. **(B)** Same format as in **(A)**, but for COCO super-categories. **(C)** Geometric properties estimated from acitivity patterns in deep neural networks for layers ordered according to relative depth (the final readout layer has relative depth of 1). All panels indicate geometric properties estimated directly from images pixels (gray violin). Note the change in scale (y-axis) across panels.

To understand effects of geometric properties on geometric SNR, we calculated average geometric properties in each ROI and plotted them in order of ascending SNR. Per the theory, for geometric SNR to increase across any transition between ROIs, at least one of the following must happen: geometric Signal increases, Dimensionality increases, or geometric Overlap decreases (assuming geometric Bias does not change). We found that no single geometric property could explain the increase in SNR across all transitions in this ordering of ROIs. However, we found that variation in geometric Signal had a strong and significant tendency to increase SNR (correlation of *ρ* = 0.48 between geometric Signal and SNR, *p <* 0.001), while variation in Dimensionality had a tendency to decrease SNR (*ρ* = −0.25, *p <* 0.05). This means that geometric Signal is the only geometric property whose variation across visual cortex has a tendency to effectively increase geometric SNR. In addition, Signal was greater in each ROI than in pixel space (gray distribution on the right in each panel of Fig. 5), while Dimensionality was smaller and Overlap was greater in each ROI than in pixel space. This means that only geometric Signal can explain why few-shot classifiers trained on brain activity patterns are more accurate than few-shot classifiers trained directly on image pixel values.

Importantly, we found that the largest Dimensionality for any concept manifold was no greater than 30, which is far less than the total number of voxels in any ROI studied here. In general, although manifold Dimensionality did show a tendency to increase with ROI size across subjects, the average number of voxels in each ROI alone could not explain average Dimensionality (Fig. S1B). For example the average Dimensionality of IPS and LO are about the same (∼ 4), but IPS has 4 times more voxels than LO (on average). Thus, variation in Dimensionality across ROIs was not a simple consequence of the differences in size between ROIs (the same goes for 5-shot classification error (Fig. S1A)).

To determine if the dependence of geometric SNR upon geometric Signal was unique to human brain activity, we investigated the profile of variation in geometric SNR for our COCO supercategories across layers of deep neural networks optimized to perform *n*-way classification of ImageNet object categories. We found that average geometric SNR increased nearly monotonically across network depth. Geometric properties varied smoothly and monotonically across early and intermediate layers, then abruptly changed direction. For example, geometric Signal decreased smoothly from the pixel layer up through 50% of the layers in AlexNet and 70% of the layers in ResNet50 before abruptly increasing across the final layers. Similar abrupt transitions were seen in the late layers for Dimensionality and Overlap. In contrast to what we observed in the brain, variation in geometric Signal was uncorrelated (*ρ* = 0.08, *p >* 0.05) with geometric SNR across early layers of the neural networks (first 50% of layers), while Dimensionality was positively correlated (*ρ* = 0.40, *p <* 0.001). Thus, the variation of manifold geometry across early and intermediate layers of neural networks was, by our particular analysis, exactly the opposite of what was observed in the brain.

We concluded that increases in geometric SNR (and, therefore, few-shot concept classification accuracy) across visual cortex depends most strongly upon increases in geometric Signal and that, while this dependence is not unique to the human brain, it is also not inevitable: neural networks achieve increases in geometric SNR across early layers using a very different strategy than that observed in our human brain data, or in the later layers of the networks themselves.

### Variation in manifold geometry is structured by strong correlations between geometric Signal, Dimensionality, and Overlap

Inspection of the profiles of varation in geometry across brain ROIs and network layers revealed that when average Signal increased, average Dimensionality tended to decrease, and average Overlap tended to increase. Similarlly, when average Dimensionality increased, average Overlap tended to decrease. These observations inspired us to investigate correlated variation in geometric properties across pairs of concepts within brain ROIs (and DNN layers) and variation in average geometric properties across brain ROIs (and DNN layers).

Perhaps most strikingly, we found that that variation in geometric Overlap across pairs of concepts with brain ROIs and DNN layers was almost perfectly explained by variation in geo-metric Signal and Dimensionality. Specifically, a log-linear transformation of geometric Signal and Dimensionality explained 97% of the variation in geometric Overlap across concept pairs for both brain and DNN manifolds (Fig. 6A).

**Figure 6:**
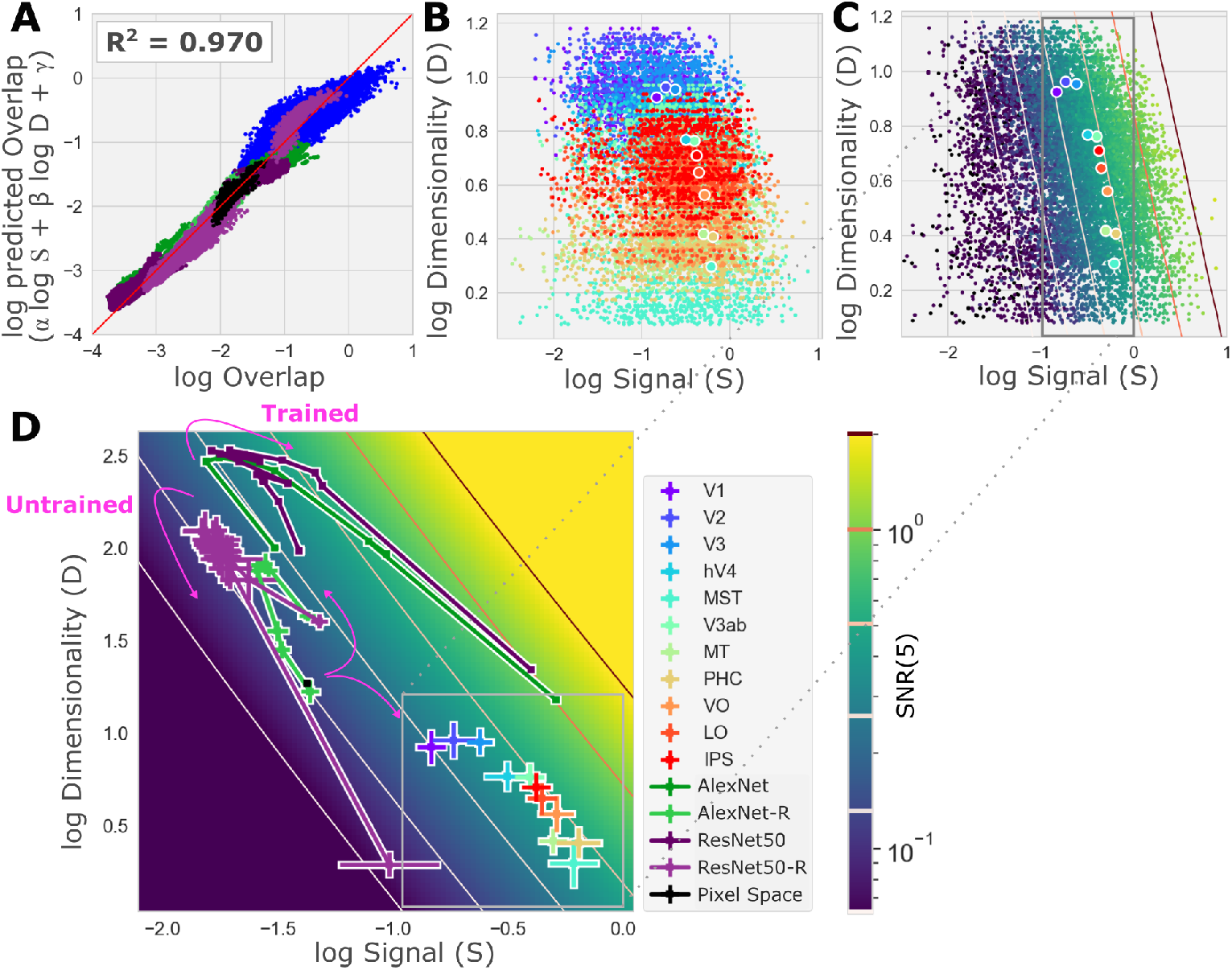
Correlations between geometric properties in brains and neural networks. **(A)** Combined Overlap predicted (y-axis) by a simple log-linear transform of Signal and Dimensionality vs. actual measured Overlap (x-axis; dots and colors as in panel **D**, all brain ROIs are blue dots). The simple model accounts for 97% of the variance (*R*^2^) in Overlap across all concept pairs, ROIs, subjects and models. **(B)** Dimensionality (*D*_*a*_, log scaled y-axis) vs. geometric Signal (*S*_*a,b*_, log scaled x-axis) for all pairs of COCO supercategories concepts (dots) for all subjects and in all ROIs (colors, panel D legend; large dots indicate ROI-wise average). **(C)** Same as panel **B** but with the concept pairs colored according to geometric SNR (color bar in panel **D**); lines of constant SNR value (“iso-SNR lines”) are drawn. **(D)** Average log Signal (x-axis) and Dimensionality (y-axis) for ROIs (box; same data as box in panel **C**) and neural network layers (“-R” indicates networks with randomized weights) on a map of SNR (colorbar). Iso-SNR lines are drawn; trained and untrained networks are labeled.

Across ROIs and layers, average geometric properties were strongly correlated. Specifically, average Dimensionality and average Overlap showed a strong negative correlation (Fig. 5) (*ρ* = −0.81, *p <* 0.001). This was an example of “coordinated” covariation, since it indicated that changes in average Dimensionality and average Overlap tended to have the same effect on average SNR. In contrast, average Dimensionality and average geometric Signal, which were strongly and significantly negatively correlated (*ρ* = −0.86, *p <* 0.001), revealed a form of “antagonistic” covariation, since increases in average Dimensionality (that would increase SNR if all other geometric properties remained fixed) tended to co-occur with decreases in average Signal (that would decrease SNR if all other geometric properties remained fixed; see Fig. S5 for a complete table of correlations between geometric properties).

To further investigate the patterns of covariation between geometric properties, we visualized the values of geometric properties in the Signal, Dimensionality (S,D)-plane. This visualization revealed quite clearly the strong negative correlation in the ROI-wise averages of geometric Signal and Dimensionality mentioned above, (larger circles in Fig. 6B), while revealing that within a single ROI, the Signal and Dimension for pairs of concept manifolds vary more independently from pair to pair (colored dots in Fig. 6B). These observations show clearly that the correlation between Dimensionality and Signal is not an inevitable relationship between geometric properties of manifolds, but rather a relationship between average geometric properties of manifolds that holds across ROIs.

The nearly deterministic relationship of Overlap to Signal and Dimensionality, and the lack of change in Bias, means that the average geometric SNR an any brain ROI and DNN layers can be computed from average geometric Signal and Dimensionality. We leveraged this finding by mapping geometric SNR across the (S,D)-plane (Figs. 6C,D).

The map of geometric SNR across the (S,D)-plane showed very clearly that the largest SNR increases occur across the earliest transitions, from pixel space to V1 and V2. There was almost no increase in SNR across intermediate areas or high-level areas (MST, V3ab, PHC, VO, LO, IPS), where variation in the (S,D)-plane was parallel to lines of fixed geometric SNR (“iso- SNR” lines in Figs. 6C,D). Thus, because of the antagonistic co-variation between Signal and Dimensionality, most of the variation in the average geometric properties of concept manifolds across higher visual cortex has little impact on classification error.

### Different profiles of geometric variation for human visual cortex and Deep Neural Networks

To visualize how the geometry of concept manifolds in human visual cortex and deep neural networks might differ, we also overlaid the average Signal and Dimensionality of each layer in several task-optimized DNNs and randomized DNNs onto the map of geometric SNR in the (S,D)-plane.

This visualization highlighted the differences in brain and DNN geometry noted earlier. The increase in SNR with the transition from pixel space (black square in Fig. 6D) to the early layers of optimized DNNs comes with an abrupt increase in Dimensionality, and no net increase in Signal. In contrast, the increase in SNR with the transition from pixel space to the early visual cortex was driven entirely by increased Signal, and occurred in spite of a decrease in Dimensionality. Interestingly, our visualizations show that after the initial increase of Dimensionality across early layers, networks abruptly switched to a geometric profile similar to what we observed in the brain: across later layers geometric SNR increases were driven entirely by increased Signal, and occurred in spite of decreased Dimensionality. Thus, the profile of geometric variation across early network layers appears to be an anomaly that distinguishes it from later layers and has no counterpart in human visual cortex.

For randomized DNNs we found that, as with trained DNNs and visual cortex, Signal and Dimensionality of pairs of concept manifolds jointly determined Overlap, while Signal and Dimensionality were negatively correlated, so that most of the variation in concept manifold geometry was along the iso-SNR lines. This suggests that the “antagonistic” covariation of average Signal and Dimensionality is a common property of both optimized and random networks. Unlike trained DNNs and visual cortex, the randomized networks did not achieve a positive net change in SNR across layers, as expected.

## Discussion

The goal of this work was to identify the geometric properties that differentiate regions of visual cortex in which concept manifolds are disentangled from regions in which they are not, and to discover any potential differences between the geometric strategies by which human visual cortex and DNNs achieve disentangled concept manifolds. Deploying the theory of Sorscher et al., we used “geometric SNR” as a proxy for disentanglement, and investigated the small set of intuitive geometric properties that collectively determine SNR.

Our strongest and mot surprising finding is that the variation of these different geometric properties across brain ROIs and DNN layers is strongly correlated. We found strong pairwise correlations between Signal, Dimensionality and Overlap, and showed that Signal and Dimensionality jointly explained most of the variation in Overlap both within and across brain ROIs (and DNN layers). The findings of strong correlation are consistent with findings in Sorscher et al., where it was shown that Dimensionality and Overlap were correlated, and that Signal and Dimensionality were correlated across DNN layers. Our work corroborates these findings and expands on them by showing that strong correlations hold for variation in geometric properties of concept manifolds across human visual cortex.

These correlations structure the variation in concept manifold geometry across brain ROIs and both trained an untrained DNN layers. Thus, they clearly do not depend on optimization with gradient descent. Although it is possible that these correlations are induced by aspects of DNN architecture that are shared with human visual cortex, we are unaware of any direct evidence for this hypothesis. The most likely hypothesis is that the correlation structure we have observed is a very general property of concept manifolds in high-dimensional spaces.

The particular pattern of correlation between geometric properties shown here has some interesting functional implications. First, as we noted above, the antagonism between Signal and Dimensionality is an example of “non-functional” covariation, in the sense that there can be a lot of variation in manifold geometry without any change in disentanglement. In the subset of high-level brain ROIs studied here, there was considerable variation in with little change in geometric SNR. It is possible that variation in geometry reflects ROI-wise difference in representation of category-orthogonal image properties (e.g., object positions, scales, poses, etc.) which were found to be increasingly decodable in concert with category properties in the ventral stream Hong et al. [2016].

Second, the negative correlation between Dimensionality and Overlap means that these geometric properties are functionally fused, in the sense that variation in each property effects change to SNR in the same direction. Given that, in addition to this constraint, covariation between average Signal and Dimensionality is antagonistic, there are effectively only two strategies that brains and networks can pursue toward disentanglement: In order to achieve significant disentanglement (increased SNR) beyond what is achievable by a random network, there must be some sequence of transitions between brain ROIs or DNN layers in which the correlation between Signal and Dimensionality is weakened such that (1) increases in geometric Signal outweigh the mitigating effect of decreasing Dimensionality, or (2) increases in Dimensionality outweigh the mitigating effect of decreasing Signal. We refer to the former strategy as the “Signal” strategy, and the latter as the “Dimension” strategy. Note that for any single transition between brain ROIs or DNN layers, these strategies are mutually exclusive.

We have provided evidence that human visual cortex follows the Signal strategy across V1– V3; afterwards the antagonistic covariation between Signal and Dimensionality dominates and, consequently, SNR barely increases. We have further provided evidence that in DNNs this Signal strategy takes effect only in the last layers. In earlier DNN layers, increases in SNR are driven by increases in Dimensionality that outweigh the mitigating decrease in Signal (the Dimension strategy). Thus, DNNs switch between the Signal and Dimension strategies.

Sorscher et al. noted that Dimensionality mediates an apparent trade-off between few-shot accuracy, which increases as Dimensionality increases, and the total number of familiar concepts that can be linearly classified by a network (Capacity), which increases as Dimensionality decreases Chung and Abbott [2021]. Thus, the Signal strategy—which is apparently pursued by the visual cortex—is implicitly capacity-increasing, as it tends to limit increase in Dimensionality; likewise, the Dimension strategy is implicitly capacity-reducing. The Signal strategy would thus appear to have an advantage over the Dimension strategy, all other things being equal.

Although it is unclear why training induces a Dimension strategy in DNNs, we note that our results are consistent with Ansuini et al. Ansuini et al. [2019], where it was shown that training DNNs induced an increase in the overall effective dimension of early layers, as well as the average number of dimensions of object manifolds, across a variety of different architectures, and using two alternate methods for estimating dimension. Dimension increase (relative to a random network) in early layers thus appears to be a very general feature of solutions to multiway classification obtained by applying gradient descent to large, overparameterized DNNs. It is also unclear why this does not seem to be the case for the brain ROIs studied here. We think it would be interesting to investigate if smaller, sparser networks that have been shown to yield the same performance as larger DNNs Frankle and Carbin [2019] would exhibit more brain-like manifold geometry in early layers.

We note an interesting discrepancy between our results and those of Sorscher et al. In their study, Dimensionality of concept manifolds encoded by spiking activity in a population of macaque V4 neurons was smaller than Dimensionality of concept manifolds in a model of V1, but that in concept Dimensionality for a population of neurons in IT was greater than in V4. They characterized this profile of geometric variation across V1,V4 and IT as “compress then expand”. In contrast, we found no evidence for this late “expansion”, and would thus describe the variation of Dimensionality of concept manifolds across human visual cortex as simply “compress”. The discrepancies between their results and ours could be due to many factors, including the differences in the species (monkey vs. human), the differences brain activity measurement (spiking activity vs. outputs of an encoding model derived from fMRI measurements), biases due to the low number of samples per concept, the circumscription of the brain regions or the fact that Sorscher et al. characterized only three regions, V4 and IT directly and V1 indirectly. We note that estimating concept manifold geometry directly on fMRI brain activity, rather than (de-noised) encoding model outputs, would not change our findings qualitatively, as it appeared to only worsen the geometric SNR and all other attendant estimates (see Fig. S4).

It is unknown if the Signal and Dimension strategies identified here will cash out in different patterns of classification decision-making, but confirming this hypothesis, and the form these different behavioral patterns might take, will require additional work. Another important aim of future work will be to deepen our understanding of how and why visual cortex controls the dimensionality of its representations. We note a tension between our results and developing evidence that low-variance dimensions that are several orders of magnitude more numerous than the Dimensionality reported throughout this work may nonetheless robustly encode stimulus information Gauthaman et al. [2024], Stringer et al. [2019].

## Materials and methods

### Data availability

The NSD dataset is freely available at http://naturalscenesdataset.org. The data are hosted in the cloud, allowing researchers to exploit high-performance cloud computing to efficiently analyze the dataset. For a complete description see Allen et al. Allen et al. [2022]. Source data are provided with this paper.

### Code availability

We provide an archive of code used in this study at https://github.com/styvesg/nsdgnet8x for the encoding model and at https://github.com/styvesg/nsd manifolds for the analysis of this paper.

### 1.1 Dataset acquisition

All data acquisition procedures were approved by the University of Minnesota Institutional Review Board.

All models were trained on the Natural Scenes Dataset (NSD Allen et al. [2022]). The NSD dataset consists of between 22K and 30K fMRI image-responses per subject (8 subjects). Images were sampled from the Common Objects in Context (COCO) database [Lin et al., 2015] and displayed at 8.4°×8.4°. The experimental design specified that each of the eight participants would view 10K distinct images (3 presentations each), and a special subset of 1K images would be shared across participants (8 subjects × 9K unique images + 1K shared images = 73K unique images); however, not all participants completed the full acquisition, so the final numbers are somewhat smaller. All fMRI data in the NSD were collected at ultra-high field (7T) using a whole-brain, 1.8-mm, 1.6-s, gradient-echo, echo-planar imaging (EPI) pulse sequence.

The image-responses are expressed in terms of betas obtained from a general linear model (GLM) analysis. For this paper, we used GLM results provided with the NSD data release, specifically, the 1.8-mm volume preparation of the data and version 3 of the GLM betas (betas fithrf GLMdenoise RR). This GLM version involves estimating the hemodynamic response function for each voxel, using the GLMdenoise technique for denoising [Kay et al., 2013], and using ridge regression to improve the estimation of single-trial betas. Betas indicate BOLD response amplitudes evoked by each stimulus trial relative to the baseline signal level present during the absence of a stimulus (“gray screen”). The betas for each voxel in each session were separately z-scored and all sessions were concatenated.

### 1.2 Identification of visual regions of interest (ROIs)s

Retinotopic visual ROIs (e.g., V1,V3,V3,V4) were identified using a separate population receptive field (pRF) retinotopic mapping experiment, as documented in [Allen et al., 2022]. Retinotopic areas (more generally described as ‘regions of interest’ (ROIs)) were manually drawn based on results of the pRF experiment. These ROIs consist of V1v, V1d, V2v, V2d, V3v, V3d, and V4, and extend from the fovea (0° eccentricity) to peripheral locations that exhibit sensi- ble responses in the pRF experiment given the limited stimulus size (the diameter of the pRF mapping stimulus was 8.4°).

Higher visual ROIs were labeled with the Kastner altas Wang et al. [2015] and provided as a reference as part of NSD. If there was any discrepancy, we gave priority to the more accurate pRF mapping labels.

Due to hardware limitation during the joint encoding model optimization, only voxels that were labeled as described above were selected. The total number of voxels (cumulative over ROI) was 11838, 10325, 11356, 9470, 9565, 11827, 9162, 10178 for subjects 1–8 respectively, totaling 83721 voxels.

### 1.3 General encoding model architecture

Brain-optimized encoding models consisted of a *shared feature extractor* (a DNN) and multiple *read-out heads* as detailed in the following sections and described in Allen et al. Allen et al. [2022] and in greater details in St-Yves et al. St-Yves et al. [2023].

### 1.3.1 Shared feature extractor

The shared feature extractors for all encoding models are sequences of transformations *e*_*L*_(*x*) = *η*_*L*_ ◦ *e*_*L−*1_(*x*) operating on *x*, here an input image, where *η*_*L*_ is the transformation that operates at layer *L* on the output of the subsequence *e*_*L−*1_(*x*). *e*_*L−*1_(*x*) and *η*_*L*_ may themselves denote arbitrary sequences of transformations. Our encoding models leverage the multiple intermediate representations *e*_*l*_(*x*), which are feature maps whose elements are denoted by [*e*_*l*_(*x*)]_*kji*_, where *k* indexes features and (*i, j*) are pixel coordinates in each feature map.

We refer to the feature extractor of the brain-optimized network as a “GNet”. The GNet feature extractor consists of a pre-filtering network (*e*_1_(*x*)) followed by a deep feature extractor. The input image resolution of the pre-filtering network is 227 × 227 and outputs an embedding of 192 features at a 27 × 27 spatial resolution. This part of the network is identical to the first two layers AlexNet.

The deep feature extractor is a sequence of blocks of layers. Each block consists of a batch normalization layer followed by a dropout layer and a convolutional layer that preserves feature map resolution. After three blocks, the resolution of the feature maps is reduced from 25 × 25 to 13 × 13 by a max pooling layer. The GNet sequence of layers was designed so that each pixel in the final convolutional layer effectively pools over the whole input image, whereas pixels in lower layers only integrate over more local regions of the input image.

#### 1.3.2 Read-out heads

Read-out heads convert features output by the feature extractor into predictions of brain activity,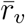, for each voxel *v*. These predictions can be expressed as a linearized model

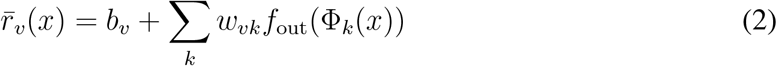

Where

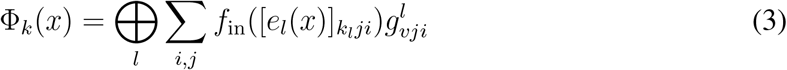

and where ⊕ denotes the concatenation along the feature axis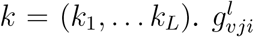 is the value of the “pooling field” for voxel *v* applied at feature maps in layer *l* at pixel (*i, j*). Pooling fields generalize the population receptive field [Dumoulin and Wandell, 2008] to arbitrary feature maps. All pooling field elements are positive-valued and normalized such that their sum equals to unity.

*f*_in_(·) and *f*_out_(·) were both set to the fully differentiable nonlinearity *f* (*x*) = tanh(*x*)log(1+ |*x*|). This nonlinearity has several interesting and desirable characteristics: 1) it has an expansive and a compressive regime, 2) it is differentiable everywhere, with no discontinuity and 3) it does not plateau over a large range.

#### 1.3.3 Objective and training

The brain-optimized encoding model was trained jointly on a large section of visually responsive cortex that totalled 83721 voxels across all 8 subjects.

The pre-filtering network was initialized from a task-optimized AlexNet and held constant for the first phase of training (see fine-tuning details in St-Yves et al. St-Yves et al. [2023]. All parameters of the deep feature extractor and the read-out heads are learned jointly via gradient descent with the ADAM optimizer (lr = 10^*−*3^, *β*_1_ = 0.9, *β*_2_ = 0.999). The training steps of the deep feature extractor and the read-out heads are alternated to promote stability of the training procedure.

To account for the noise profile and the discrepancies in voxel predictability, we used the following *L*2-norm weighted loss function

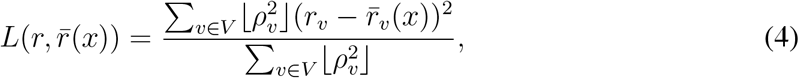

where 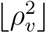 is the batchwise Pearson-like correlation of every voxel with a floor of 0.1 to always permit some contribution of the yet-to-be-predicted voxels (i.e. those with low correlation at the onset of training) in the voxel ensemble *V*. If the model can be trained voxelwise, then this weighting could be ignored but this could affect the learning dynamics (learning rates).

Further details regarding the regularization procedure and fine-tuning can be found in StYves et al. St-Yves et al. [2023].

### 1.4 Details on concept manifold geometry calculation

To evaluate concept manifold geometry under the theory of Sorscher et al. Sorscher et al. [2022] we adapted code provided with the original manuscript. Briefly, we represent every set of activity patterns (fMRI, encoding model, or neural network) corresponding to *P* examplars from a concept *a* in a *V* -dimensional feature space as a *P* -by-*V* matrix *X*^*a*^. We calculate the centroid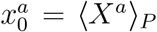 and perform principal component analysis to calculate the variance explained *a*_*i*_ along the principal component direction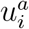 for *i* ∈ [0, *V* − 1]. Likewise for any other concept *b*.

From there, we calculate the radii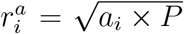, average concept radius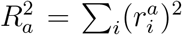,normalized pairwise centroid distance 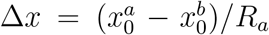 with respect to concept *a*, and normalized direction matrices

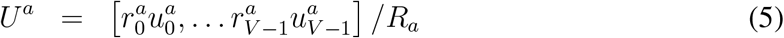

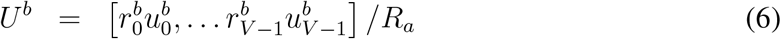

from which all other terms involved in Eq. 1 are expressed. The geometric Signal and Bias are given by S_*a,b*_ = ||Δ*x*||^2^ and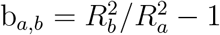, respectively. The concept-wise Dimensionality is defined by the participation ratio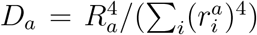. The geometric Overlap terms are given by 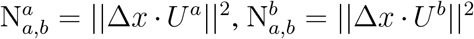 and 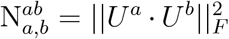where || · ||_*F*_ is the Frobenius matrix norm, which is the sum of the square of every entry. We did not consider any terms in the theory involving power-of-*m* greater than 1, where *m* is the number of examplars defining the concept prototype under *m*-shot learning.

### 1.5 Datasets, geometry and empirical few-shot accuracy

We initially performed the geometry estimation in the native voxel space and found that the highest participation ratio dimension was of the order of 20–30. Based on this result, and to facilitate future geometry estimation, we followed Sorscher et al. [2022] and randomly projected all ROI voxel populations into a 300-dimensions random subspace. Given the relatively large number of dimensions of this subspace w.r.t. the dimensions of interest, we found that all geometry estimates were accurately recovered as predicted by the theory (see Fig. S3).

Once the encoding model had been trained, we compared geometric estimates derived from synthetic brain activity patterns output by our encoding model to estimates derived from directly measured brain activity (which was used to train the model, incidentally). For this analysis, visual concepts were the 12 ‘thing’ super-categories of the COCO dataset (Rev. 2017). The COCO super-categories are: ‘appliance’, ‘accessory’, ‘kitchen’, ‘person’, ‘indoor’, ‘outdoor’, ‘vehicle’, ‘electronic’, ‘sports’, ‘food’, ‘furniture’ and ‘animal’. Each of these supercategories contains several constituent COCO sub-categories. For example, the ‘outdoor’ supercategory includes the sub-categories ‘traffic light’, ‘fire hydrant’, ‘stop sign’, ‘parking meter’ and ‘bench’. These are only loosely associated, and it should be understood that the supercategory ‘outdoor’ here stands for this collection of constituents, not to a broader concept of ‘outdoor’ (e.g., like whether the image depicts an outdoor scene). This is not an issue for our analysis since the concepts are arbitrary and only serves as probes for the representational geometry. For more details, see Fig. S2 or refer to the 2017 COCO things annotation files found at https://cocodataset.org/. Note that images in the COCO dataset may contain multiple objects corresponding to different super-categories. If any of pixel in a COCO image (cropped as described in Allen et al. Allen et al. [2022]) depicted a super-category, it was labeled as belonging to that super-category. Therefore, any one image in the COCO dataset may contributed to multiple concept manifolds. This explains the decrease of geometric SNR of COCO supercategory concept manifolds relative to ImageNet concept manifolds.

When comparing geometric estimates derived from encoding models and direct brain activity measurements, we did not need to use the same training/validation dataset split used for training of the encoding model, since the question here is not one of cross-validation but of characterization. However, we did need to validate the geometric theory. In order to do so, we performed multiple folds of geometry estimation and empirical few-shot error calculation. For each subject, the roughly 10k image responses are randomized and divided into two groups of roughly 90% and 10% of samples. The geometry is calculated on each concept member in the larger group and the smaller group is randomized and further divided into a group of *m* sample per concept and those remaining. The *m*-shot prototype is calculated and used to classify the remaining samples of the smaller group according to the few-shot accuracy classification rule. This latter randomization is performed 16 times and the result is averaged. The larger randomization loop is also performed 16 times and the average and standard deviation of the geometric characteristic, SNR and empirical *m*-shot accuracy is calculated. Fig. 4 shows the average empirical 5-shot accuracy plotted against the average SNR calculated in this manner. Fig. 5 shows the average geometric SNR components: the Signal, Bias, noise-signal Overlap and noise-noise Overlap calculated in this manner, averaged over pairs of concepts.

We also estimated geometric properties for concept manifolds corresponding to cthe 1,000 classes of the ImageNet 2012 database Deng et al. [2009]. Note that this was only possible using an encoding model, as ImageNet stimuli were not included in the NSD. For this analysis, we kept the original training and validation image separation. The geometry was calculated over the larger training set samples for each category. We randomly selected *P* = 700 images from each category to estimate the geometry. The ImageNet validation set was used to calculate a multifold 5-shot accuracy in the manner described above, with the number of folds set to 16. We also performed the geometry estimation in the original space in each ROI. The results highlighted the invariance of the geometry estimates under sufficiently high dimensional random projections.

### 1.6 Extracting layerwise DNN concept geometry

We looked at two networks’ internal representations at various stages along the network hierarchy. In most cases, We extracted network activities after the activation function following either a convolution or fully connected layer, which occurs in any network at relatively regular intervals. These internal representations (features) were randomly sampled into a lower dimensional space of 50K dimensions (or the full space if the dimension was lower) and then randomly projected onto 2,000 dimensions. This was done in order to reduce the computational (and memory) cost of the random projection for very large layers. Comparison with and without this random projection in case where not applying it could be afforded showed extremely comparable values for all the geometric estimates. Likewise, the choice of the number of projection dimension was such that all estimates were sufficiently close to convergence. In all cases, stability of the relevant estimates was the deciding factor for any embedding dimension.

All models parameters were imported from the pyTorch (v1.10.0+cu113) torchvision (v0.11.1) model library. For all models, we performed the analysis of concept geometry evoked by the NSD COCO images with the trained model weights (trained) and with a randomized version of the trained weights where all network parameters have been shuffled (untrained).

The geometry estimation was performed 9 times for each extracted layers with a new shuffling of the weights for the untrained variant, and different randomly sampled features and random projection each time the analysis was performed. The values plotted in Figure 5 and Figure 6 show the mean and std. dev. over these 9 estimates. The “trained” model had std. dev. on their estimate of Dimensionality and Signal smaller than the symbol size (Fig. 6), indicating the high degree of consistency of these geometric properties w.r.t the feature sampling procedure. Therefore, the sampling step that introduced the greatest source of variance in this procedure appears to be the shuffling of the parameters in the “untrained” model estimate. All other sampling offered very robust estimates.

### 1.7 Combined geometric Overlap term

We initially computed signal-noise Overlap and noise-noise Overlap as defined by Sorscher et al. Sorscher et al. [2022]. However, in all network and brain manifolds, these properties were highly correlated. Therefore, in Figure 5 we described the combined geometric Overlap term

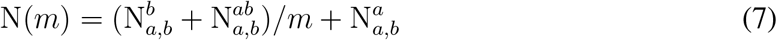

and later in Figure 6 we predicted this term as a function of Signal and Dimension. Note that this combined term depends on the number of examplar *m* in the concept prototype. Therefore, the regression parameters will be dependent on *m* since Signal and Dimensionality are independent of *m*.

### 1.8 Relating Signal, Dimensionality, and Overlap

After performing principal component analysis on the space of representational geometry Signal, Bias, Dimensionality and combined Overlap), we observed empirically that two factors accounted for most of the variance (greater than 92% of variance) in all the representations that we measured (brains and DNNs). These factors are in the plane of Signal and Dimensionality which implies that all other properties can be relatively accurately described as a linear combination of these two properties. The Bias term being essentially independent and constant, this leaves only the combined Overlap term which we write as

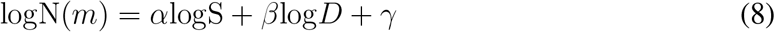

A 2-dimensional linear regression of this functional form for all model estimate in Figure 6A jointly yielded a coefficient of determination *R*^2^ = 0.970. By comparison, a linear regression of the combined Overlap term onto the Signal alone produced a correlation of 0.56.

The fitted values of (*α, β, γ*) enable one to write SNR as a function of Signal and Dimensionality only. We indicate SNR using using color gradients and isolines in Figure 6A. Note that the isolines are slightly curved in log-log space. The regression can be performed independently for concept manifolds in brain ROIs and DNN layers indepedently, or as a whole. While there are some deviations from one model to another, the results are largely consistent; this finding warrants a comparison of the models in a single common (*S, D*)-plane as shown in Figure 6D.

Figure 6C shows that the relationship between Signal, Dimensionality and SNR also holds for individual concept pairs. Concept pairs whose geometry has been estimated in all brain ROIs have been combined in this plot which shows that the 2D determination is obeyed at the concept level in all ROIs simultaneously.

### 1.9 Symmetry of 5-shot error matrices

Fig. 3C shows the correlation between the ImageNet hierarchical distance matrix and the fewshot success rate matrices. The ImageNet hierarchical distance matrix is always symmetric, whereas the few-shot accuracy matrix need not be. To measure the symmetry of few-shot error matrices, we calculated the correlation between the lower and upper triangular component of the few-shot accuracy matrices (ROI-wise). The result is shown in the inset of Fig. 3C.

### 1.10 Correlations of SNR and SNR components

While the geometrical SNR is determined by Eq. 1, the geometric Signal, Bias, Overlap and Dimensionality all depends on common factors which induces correlations in these quantities, therefore a simple look at the dependence of geometric SNR on its components does not reveal the net influence that changing these properties have on geometric SNR.

Therefore, we calculated the empirical Pearson correlations between the geometric components and SNR for the mean of ROI (layer for networks) estimate and for individual concept manifold pairs, for all ROIs (layers) and subjects (samples) concatenated together (referred to in the text and shown in Fig. S5). For the networks, we concatenated layers from AlexNet and ResNet50 together in a single estimate since both showed similar trajectories. Significance was estimated by performing 1,000 two-sided permutation test.

## Acknowledgements

This work was supported by NSF CRCNS grant IIS-1822929 and NIH grant R01EY023384 (T.N.) and NIH grant R01EY034118 (K.K.).

## Author contributions

G.S.-Y. and T.N. conceived of the project and wrote the paper. G.S.-Y. performed data analysis. G.S.-Y., K.K, T.N. directed the overall project. All authors discussed and edited the paper.

