## Supplementary Information for "Variation in the geometry of concept manifolds across human visual cortex"

### Supplementary figures

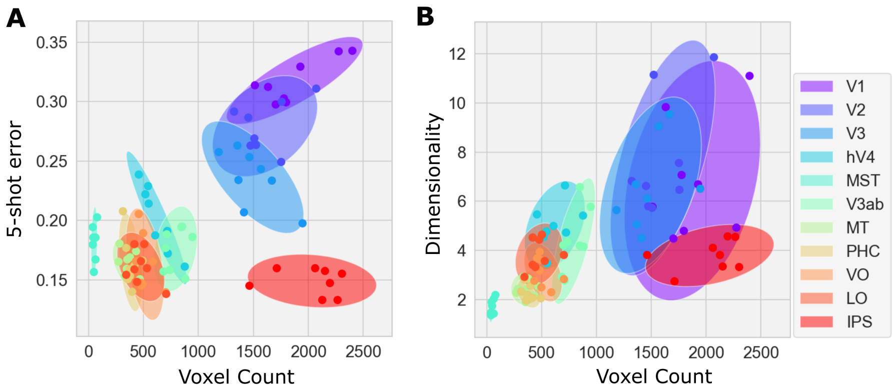


**Supplementary Figure 1: Relation of A) few-shot accuracy and B) Dimensionality to ROI voxel count.** The point correspond to the estimate of the 8 subjects and the ellipses capture 2 std. dev. around the subject average. There is a tendency for ROI higher in the visual hierarchy to have smaller number of constituent voxels. Could this fact alone explain the change in few-shot accuracy and dimensionality? It appears this is not the case since at least one higher ROI, IPS, has constituent voxel count on the same order as early visual areas but nevertheless has vastly superior (inferior) few-shot accuracy (Dimensionality).

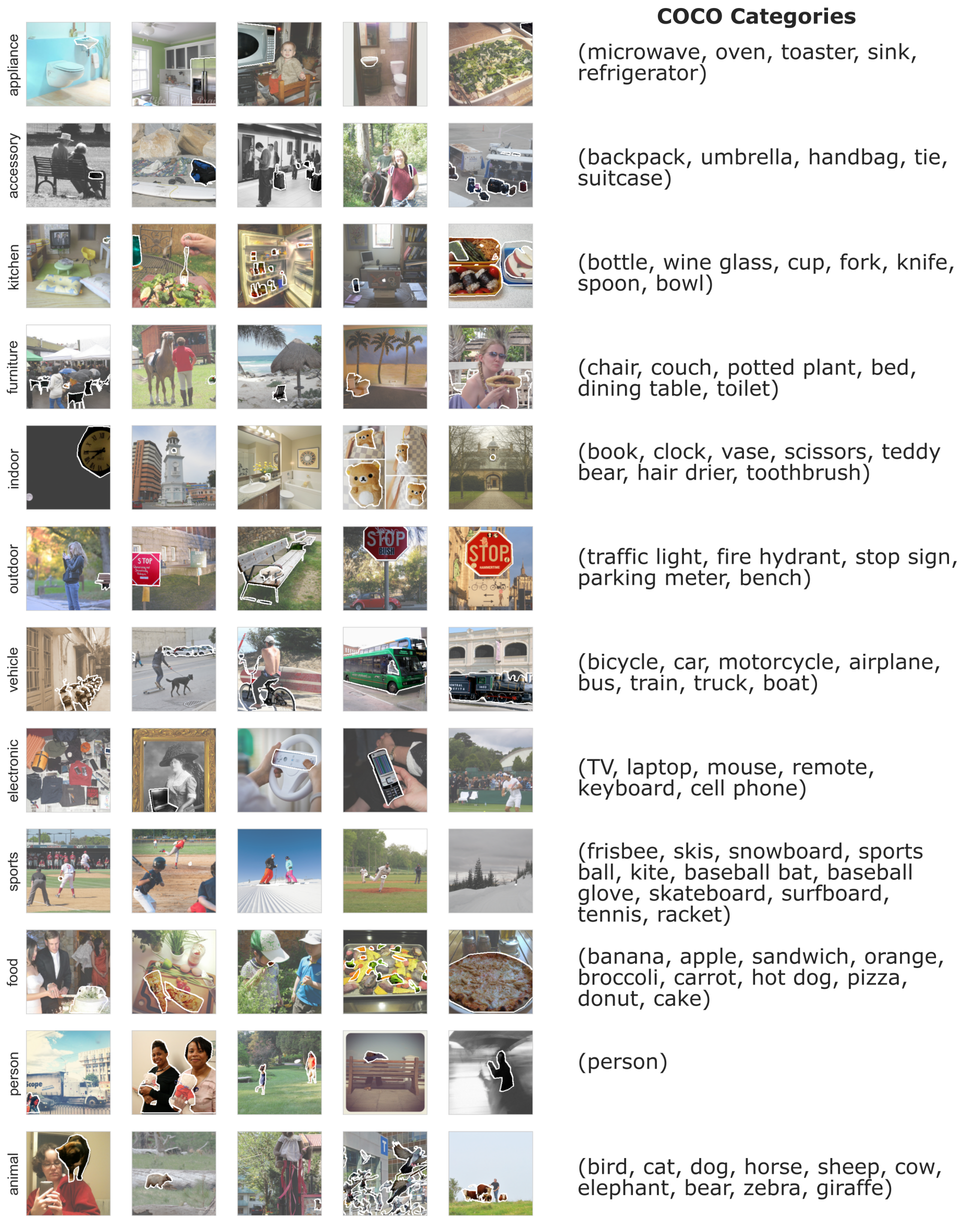


**Supplementary Figure 2: Example images from each COCO supercategories used to probe concept manifolds. A)** An image is deemed an examplar of a supercategory manifold if some of its pixels belong to an object of one of its constituent objects. The object belonging to the supercategory have been highlighted with a white surrounding and the remaining pixels have been shaded out slightly to emphasize the pixels determining the supercategory attribution **B)** The list of the constituent objects for each supercategory.

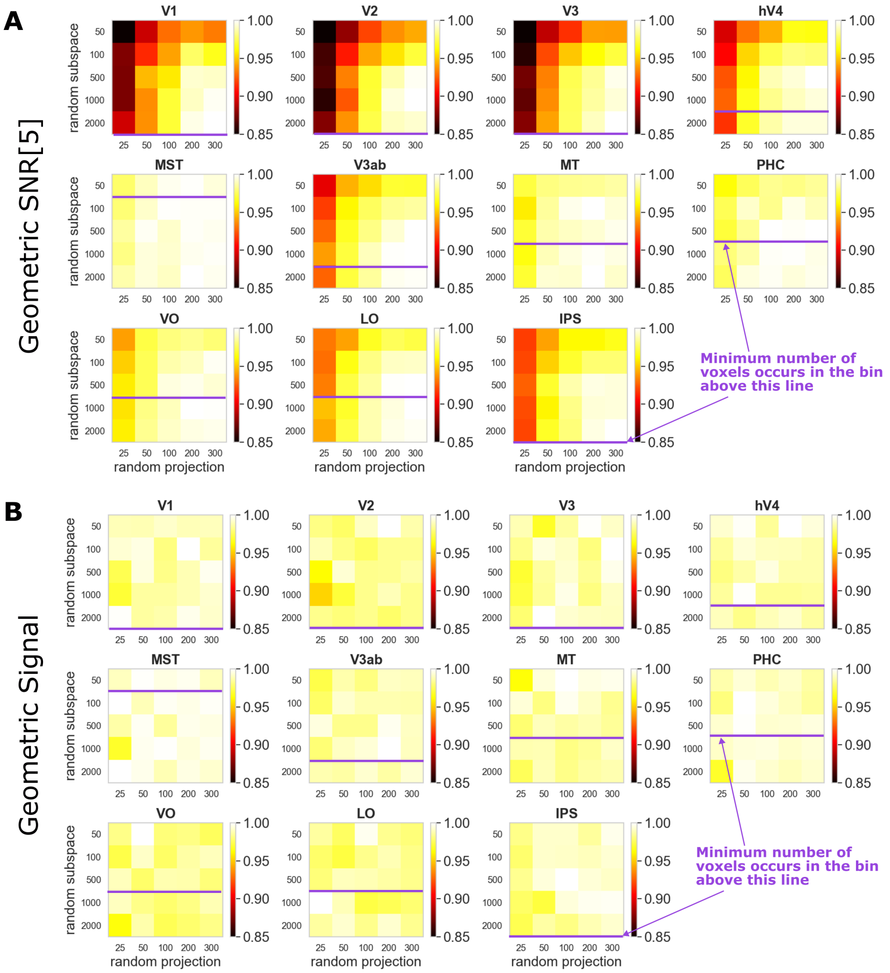


**Supplementary Figure 3: Convergence of the geometric estimates.** We show the convergence of the geometric estimate as a function of two of the representation embedding hyperparameters: the random subspace dimension (i.e., the number of randomly selected voxels) and the random projection dimension (i.e., the number of random dimensions onto which activity patterns are projected). **A)** Each panel show the fraction of the estimate of Geometric SNR[5] relative to its maximum in various regions. **B)** Each panel show the fraction of the estimate of Geometric Signal relative to its maximum in various regions. In both cases, the purple lines provide an estimate of where the random subspace dimension become greater than the number of voxels in that ROI. If that dimension is greater than the number of voxels, then all voxels are always selected. In the main figures of the paper, we used all available voxels in each brain ROI; for networks, the random subspace dimension was fixed at 25,000.

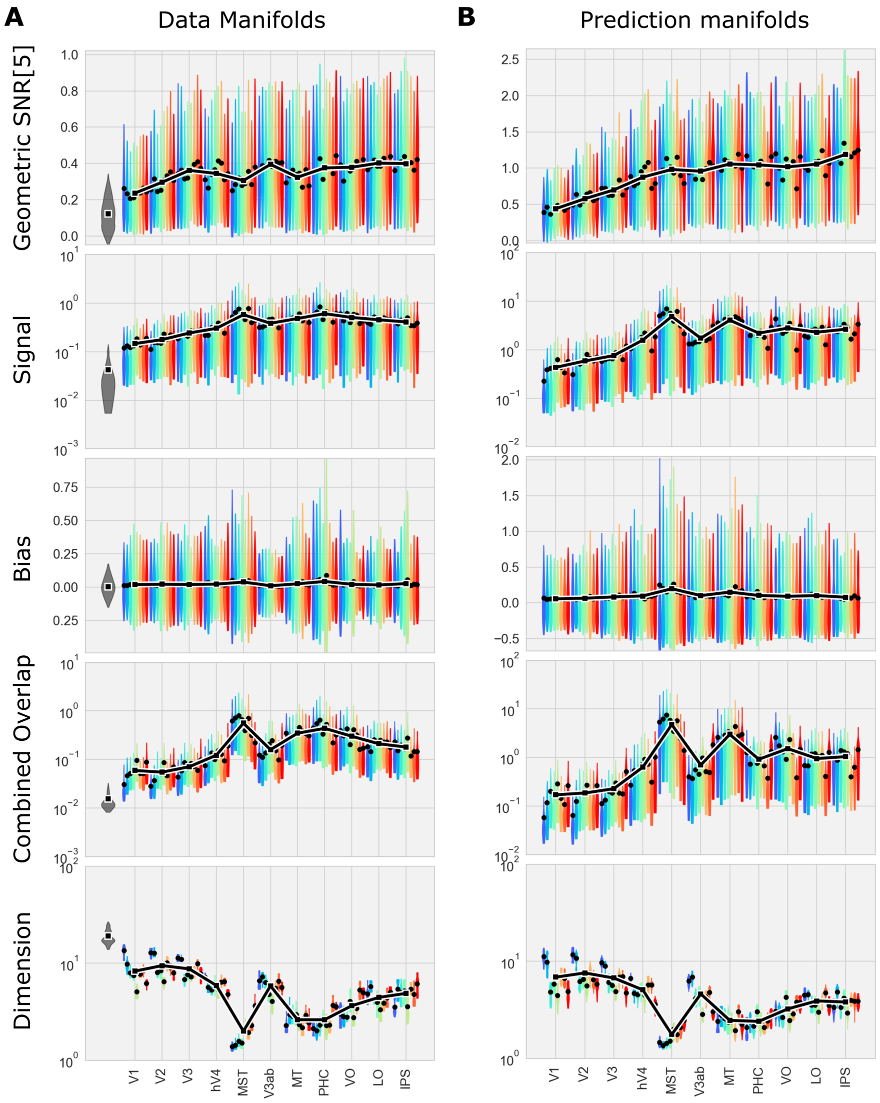


**Supplementary Figure 4: Comparison of the geometry estimate on A) measured fMRI data and B) on predicted fMRI data.** We notice the overall reduction in SNR, Signal and, counter-intuitively, Overlap, while Dimensionality increases for manifold geometry estimated directly on data. On the other hand, the overall pattern appears very similar, with the same antagonistic variation in geometric components.

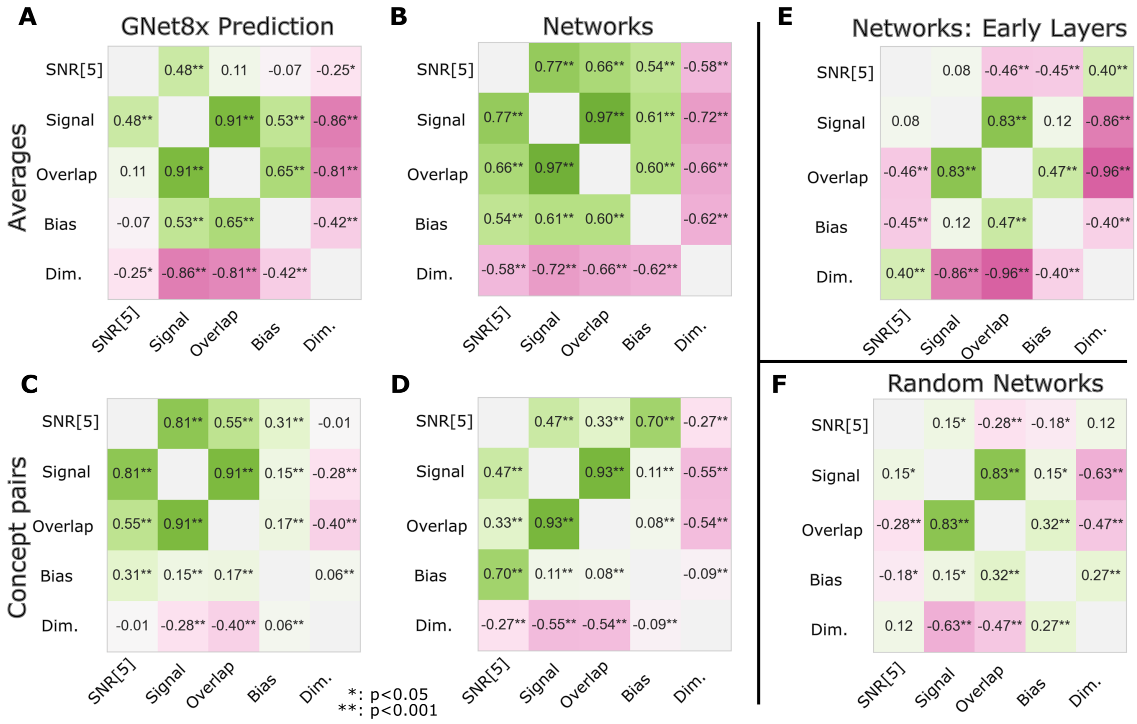


**Supplementary Figure 5: Empirical correlations between the geometric components.** Correlations of the average of the estimates for **A)** brains **B)** and networks. Correlations for all concept pairs of the estimates for **C)** brains **D)** and networks. **E)** Correlation of the average of the estimates for early layers (layers with fractional depth smaller than 0.5) only. **F)** Correlation of the average of the estimates for randomized networks.
